# Genetic Architecture of Perivascular Space Morphology in the Pediatric Brain

**DOI:** 10.64898/2026.05.21.726913

**Authors:** Jessica Morrel, Hedyeh Ahmadi, Carinna Torgerson, Rachel Custer, Haoyu Lan, W. James Gauderman, Jeiran Choupan, Megan M. Herting

## Abstract

**Introduction:** Perivascular spaces (PVS) support brain homeostasis through metabolite delivery and waste clearance, yet the genetic determinants of PVS morphology during childhood remain unknown. Here, we leveraged cross-sectional Adolescent Brain Cognitive Development Study data (N = 6,600; ages 9-10), including genomics and 3T structural MRI.

**Methods:** Linear mixed-effects models examined associations between 45 single nucleotide polymorphisms (SNPs) previously linked to adult PVS structure or function and PVS count and volume fraction (VF) across six macroregions and 28 Desikan-Killiany subregions.

**Results:** Fifteen SNPs demonstrated significant associations with PVS macroregion morphology, predominantly VF; 21 SNPs demonstrated associations with subregion morphology. Variants near *SLC13A3* showed the strongest, most widespread associations with PVS VF. Additional replicated variants implicated Wnt signaling, cell adhesion, apoptosis, and glymphatic function.

**Conclusion:** These findings suggest genetic associations with PVS morphology are detectable in childhood, while highlighting developmental and regional specificity. Longitudinal studies are now needed to determine whether childhood PVS genetic architecture predicts trajectories of glymphatic maturation and associated cognitive outcomes.

**Key Points:**

- Perivascular space morphology, mapped across 28 white matter subregions, displays substantial regional and interindividual variability during childhood.
- Six genetic associations with perivascular morphology were only detectable at subregion resolution, demonstrating that lobar averaging can obscure spatially restricted signals.
- Nearly 50% of adult-identified perivascular space risk variants replicated in 9-10 year-olds, with anatomically patterned associations reflecting patterns of neurovascular complexity and neurodevelopmental timing.

## 1. Introduction

Childhood is a critical period of neurodevelopment characterized by rapid (*1*) and highly metabolically costly (*2*) changes to brain structure and function. As such, efficient nutrient delivery and waste clearance are key to maintaining brain health and supporting neural maturation; however, the biological systems subserving these functions during childhood remain poorly understood. Perivascular spaces (PVS), as part of the brain clearance system, may represent one such critical system that supports these metabolic demands. PVS are cerebrospinal fluid (CSF)-filled spaces surrounding cerebral blood vessels extending into the brain (*3–5*). Working alongside the blood-brain barrier (BBB), PVS help maintain homeostasis (*6*) and support waste clearance, solute delivery, and CSF-interstitial fluid exchange (*7, 8*). While human and animal studies have highlighted the role of PVS in neurological health and disease (*9–17*), our understanding of PVS morphology, and the factors that shape it, during development remains limited (*16, 18*), as the current body of literature has focused on PVS burden during adulthood and neurodegeneration (*11–13, 17*). Recent advancements in magnetic resonance imaging (MRI) and PVS segmentation techniques (*19–23*) have allowed for mapping of smaller PVS across the brain, enabling their application in healthy and developing populations. Burgeoning evidence suggests that MRI-visible PVS are present in healthy neonates (*24*), children (*25–31*), and adolescents (*25–29*). Notably, PVS count and volume fraction (VF) exhibit marked interindividual variability and a regionally specific distribution across the brain in children and adolescents (*27, 28*). Whether genetic factors contribute to individual- and regional-level differences in PVS structure during this critical developmental window, however, has yet to be established.

Several adult studies have provided compelling evidence supporting a relationship between genetics and PVS (*32–35*). In a sample of approximately 900 young adults from the Human Connectome Project (HCP), PVS to white matter ratios were shown to be significantly more similar in siblings, and particularly, monozygotic twins compared to non-siblings, suggesting a genetic component to PVS burden (*35*). Building on this finding, a recent genome-wide association study (GWAS) on over 40,000 adults identified 24 genome-wide loci that were significantly associated with higher PVS burden (number of enlarged PVS) in the white matter, basal ganglia, and hippocampus, with white matter PVS displaying the highest degree of heritability (*32*). The identified loci implicate pathways involved in cerebrovascular function, including blood vessel diameter, BBB integrity and function, expression of tight junction proteins and aquaporin-4 (AQP4), and angiogenesis (*32*). AQP4 plays a central mechanistic role in glymphatic flow, perivascular fluid transport, and brain–CSF exchange (*36, 37*). Genetic variation in the *AQP4* gene moderated the relationship between sleep quality and duration and Aβ-amyloid burden, implicating its expression in sleep-dependent glymphatic function (*34*). Beyond PVS burden, a recent GWAS of 40,000 European adults from the UK Biobank identified 17 single nucleotide polymorphisms (SNPs) associated with diffusion tensor imaging along the PVS (i.e., DTI-ALPS; (*38*)) which, although limited, has been proposed as an indirect measure of glymphatic system function (*33*). DTI-ALPS associated loci implicated a number of nervous system processes, such as dopaminergic signaling and dendritic growth, with no overlap with the aforementioned loci (*33, 34*). Notably, the loci associated with PVS burden replicated in young adults (*32*), and a subset are enriched in genes linked to early-onset leukodystrophy, a condition affecting pediatric white matter, and expressed in fetal brain endothelial cells (e.g., *GFAP, SLC13A3, PNPT1*). Yet, no study has examined whether these genetic factors shape PVS morphology during childhood, when the neurovascular system is still maturing, or whether the same risk variants associated with PVS burden in adults exert detectable effects earlier in development, before the accumulation of vascular risk factors.

To address these gaps, we leverage the Adolescent Brain Cognitive Development (ABCD) Study, a large, diverse cohort with harmonized MRI acquisition protocols and genetic sequencing, to provide the first examination of genetic influences on PVS morphology during childhood. Using a high-resolution PVS segmentation pipeline, we test whether SNPs identified through the GWAS of adult PVS burden (*32*) and DTI-ALPS (*33*), and AQP4 variants implicated in glymphatic flow and brain-CSF homeostasis (*34*), are associated with PVS count and volume fraction (VF) at ages 9-10. Count models assess whether genetic variation relates to having a greater number of MRI-visible PVS, whereas VF models evaluate overall PVS burden, capturing dilation independent of count. Given that PVS prevalence and morphology vary substantially across white matter regions in children and adolescents (*27*), we examined genetic associations at both macroregion and subregion levels of spatial resolution, allowing us to determine whether genetic effects are broad or anatomically specific. We hypothesize that SNPs near genes involved in astrocyte structure and function, regulating cellular migration, and angiogenesis will be associated with PVS burden in the frontal and parietal white matter, as PVS dilations in the white matter of these regions has been shown to be most prevalent at this developmental stage (*15*).

## 2. Materials and Methods

### 2.1 Study Design

This study used cross-sectional imaging and genetic data collected at the baseline visit of the Adolescent Brain Cognitive Development^SM^ (ABCD) Study^®^ (NIMH Data Archive annual 3.0 [raw T1 and T2 images] and 5.0 [all other data] releases; DOI: 10.15154/8873-zj6, DOI: 10.15154/1520591). In brief, the ABCD Study enrolled over 11,800 9-10-year-olds (mean age = 9.49 years; 48% female) between October 2016 and October 2018 from 21 sites across the United States (*64, 65*). Centralized Institutional Review Board (IRB) approval was obtained from the University of California San Diego and each site obtained approval from its respective local IRB. Each child’s parent or legal guardian provided written consent, and each child provided written assent. Eligibility requirements for the study included 9.0-10.99 years at the baseline visit and English fluency. Participants were excluded if they had severe sensory, neurological, medical, or intellectual limitations, or were unable to complete the MRI scan. ABCD participants were included in this study if they had genetic data at baseline and passed all neuroimaging and PVS segmentation quality control (QC) criteria (**Section 2.3**). To ensure independence of observations, analyses were restricted to one randomly selected child per family, resulting in 6,600 participants in the current study sample (see **Figure S1** for flowchart of participant exclusion).

### 2.2 Genotyping and Quality Control

Genomic data for the ABCD Study were generated from saliva and blood samples. Specimen storage, DNA extraction, and genotyping calls were performed at the Rutgers University Cell and DNA Repository (RUCDR) as previously described (*66*). Briefly, DNA extraction and genotyping were performed using the Affymetrix Smokescreen array (*67*) resulting in over 700,000 single nucleotide polymorphisms (SNPs). After QC, imputation was performed using the TOPMED reference panel (*68*), resulting in over 307 million imputed variants available using reference GRCh38. Population structure was characterized using PC-AiR from the GENESIS package resulting in 32 ancestry principal components (*69*). SNPs with a call rate <98% and/or a minor allele frequency <5%, and individuals with extreme heterozygosity, were excluded. Of the 52 previously identified SNPs, 45 were successfully extracted from our dataset (**Table 1**, see **Table S1** for gene and function mapping); the remaining 7 SNPs did not pass genetic QC. Genotype data were converted to PLINK format, and the 45 SNPs of interest were extracted (**Tables 1, S1**). The source paper, or, when not present, the Ensembl Variant Effect Predictor (VEP) tool was used to estimate SNP consequences and gene mappings (*70*).

**Table 1.**
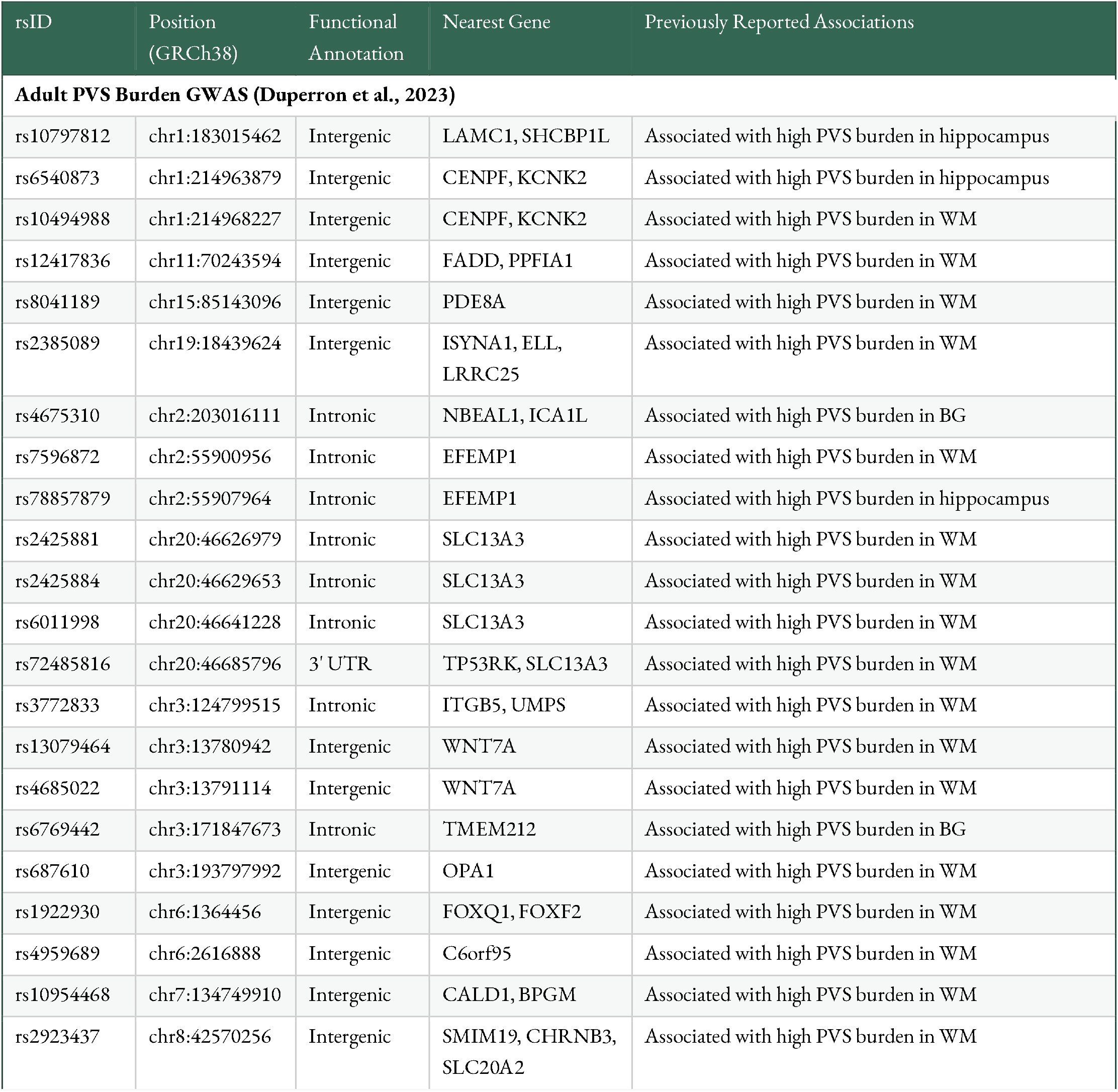
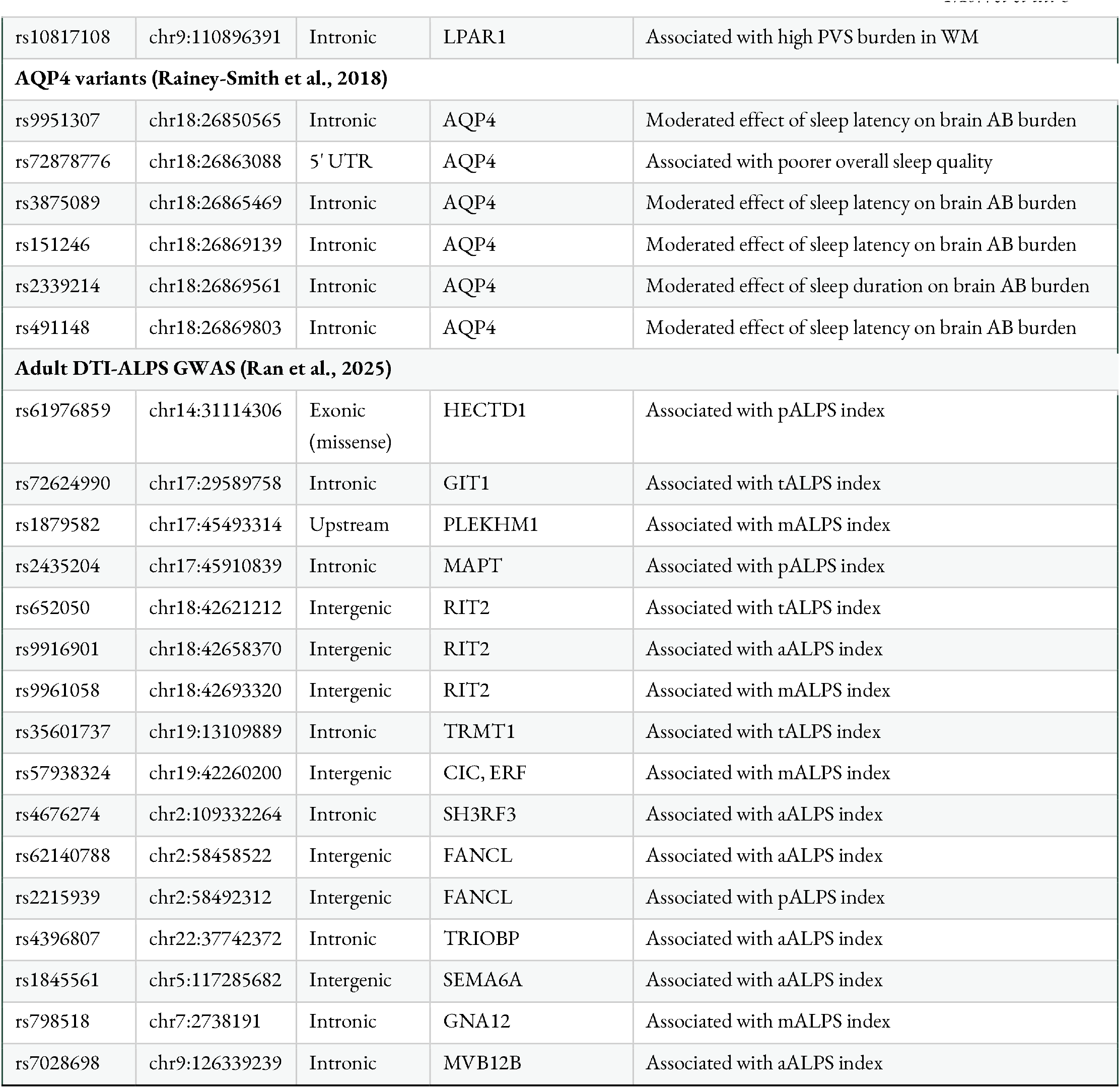
Single nucleotide polymorphisms (SNPs) previously associated with perivascular space (PVS) structure, diffusion, or glymphatic function in adults. SNPs presented along with their functional annotation (i.e., most severe consequence) and nearest gene as well as the brain outcome each SNP was previously associated with in adults. Abbreviations: Basal Ganglia (BG), White Matter (WM), Beta Amyloid (AB), middle (mALPS), posterior (pALPS), anterior (aALPS), total (tALPS) diffusion tensor imaging along the perivascular space (DTI-ALPS).

### 2.3 Neuroimaging Data

Given the susceptibility of PVS morphology to head motion and imaging artifacts, we restricted analyses to participants whose raw T1- and T2-weighted images met ABCD Study’s QC criteria, including clinical findings, signal-to-noise ratio, artifact, and motion (*45*; **Figure S1**). Raw T1 and T2-weighted images were subsequently processed using the Human Connectome Project (HCP) minimal preprocessing pipeline (*72*), including upsampling to 0.7-mm isotropic resolution to improve tissue contrast. Following preprocessing, T1w and T2w images were denoised using adaptive non-local means filtering (*73*), and enhanced PVS contrast (EPC) images were generated by computing the ratio of filtered T1w to T2w images to improve PVS contrast visibility (*22*). All EPC images underwent manual visual inspection, and participants exhibiting excessive noise or incidental findings were excluded. PVS segmentation was then performed using the Weakly Supervised Perivascular Spaces Segmentation framework, as previously described (*74*). White matter was parcellated according to the Desikan–Killiany (DK) atlas (subregions; *49*), yielding both 34 bilateral subregion-level parcellations and six macroregions: frontal lobe, temporal lobe, parietal lobe, occipital lobe, the cingulate, and centrum semiovale (CSO) (see **Table S2** for areas comprising CSO) of the white matter. The insula was grouped within the cingulate macroregion given its anatomical proximity. Within each spatial unit of analysis (i.e., DK subregion and six macroregions), PVS burden was quantified using both count and volume fractions (VF), the latter defined as the ratio of PVS volume to regional volume (i.e., CSO PVS volume/CSO total volume), normalizing for anatomical size differences between subjects for each region of interest. Given that PVS prevalence and morphology have been found to vary substantially across white matter regions in children and adolescents (*27*), we examined genetic associations at both levels of spatial resolution. Macroregion analyses, focused on each lobe, the cingulate, and the CSO, provided broad characterization of genetic effects, while subregion analyses within each lobe allowed us to determine whether genetic associations were regionally specific or uniform within each lobe, a distinction with implications for understanding the anatomical specificity of genetic influences on PVS structure.

### 2.4 Demographics & Covariates

Average household income in US Dollars (i.e., *<50K*, ≥*50 and <100K*, ≥*100K, Don’t Know/Refuse to Answer*), age at MRI (in months), and sex (*female* or *male*) were included due to known relationships with PVS structure and function (*26, 27, 33, 35*). Genetic ancestry principal components (PCs) 1–5 were determined to well-capture the genetic variation of the study sample (**Figure S2**) and were therefore included in all analyses to account for population stratification and differences in brain morphology across ancestries (*76*), as continuous genetic ancestry captures admixture more precisely than self-reported race/ethnicity (*77*). All models also included a variable to simultaneously capture MRI model and headcoil (e.g., 32-channel Siemens Prisma) to account for differences in PVS due to variability in scanner differences. ABCD site was included as a random effect to account for the hierarchical nature of the ABCD Study. Lastly, to account for differences in regional volume when modeling PVS counts, total white matter volume of the region of interest was included as a covariate; this correction was not necessary for VF models, as volume fraction is calculated relative to regional volume and thus inherently controls for anatomical size differences.

### 2.5 Statistical Analyses

In primary analyses, linear mixed effects (LME) models were implemented to test the independent associations between each of the 45 preselected SNPs and count or volume fraction (VF) of each macroregion (6 regions x 2 metrics; N = 12 outcomes) (see **Supplemental Methods 1.1** for model equations). Analyses using count as the outcome evaluate whether genetics are related to having more identifiable PVS, suggesting a higher frequency of MRI-visible spaces. VF analyses examine whether genetics are associated with a greater overall PVS burden in a region. When interpreted alongside count models, elevated VF in the absence of higher counts would suggest that individual PVS are larger or more dilated, whereas concordant effects across both metrics would implicate greater PVS number as the primary driver. Occipital VF was log-transformed prior to analysis to satisfy linear regression assumptions (see **Supplemental Methods 1.2**). To control for multiple comparisons while preserving power to detect region- and metric-specific effects, false discovery rate (FDR) correction using the Benjamini–Hochberg procedure was applied separately within each region-metric combination (i.e., 45 tests per correction, 12 corrections total). Results with FDR-adjusted P < 0.05 were considered statistically significant.

To determine whether genetic associations with PVS burden were uniform across a macroregion or driven by specific subregions, we conducted follow-up LME analyses using 28 bilateral DK atlas subregions grouped into the macroregions they belong to: frontal (9 subregions), temporal (5 subregions), parietal (5 subregions), occipital (4 subregions), and cingulate (5 subregions) (**Table S2**). Specifically, for each of the 45 SNPs, models were conducted for count or VF, with DK subregion nested within each macroregion as a predictor interacting with SNP (see **Supplemental Methods 1.3** for model equations and **Supplemental Methods 1.2** for related model diagnostics). Region main effects were excluded from models because outcome standardization (which was necessary to satisfy modeling assumptions) centered each region at zero, absorbing region-level mean variation and rendering region intercepts uninterpretable. Consequently, the models capture within-macroregion variation across subregions, rather than between-region differences, and specifically test whether genetic effects differ across subregions independent of overall regional mean differences. The same covariates as the macroregion models were retained, including regional volume as an additional covariate in count models. In addition to a random intercept for site, these models also included a random intercept for participants as nested subregions are within the same individual. For each model, the SNP x subregion interaction was tested using Type III ANOVA with Satterthwaite approximation for degrees of, evaluating whether the genetic effect on PVS burden varied significantly across subregions within each macroregion. For each model, we implemented estimated marginal trends to estimate the SNP effect within each DK subregion, while holding covariates at their reference values. The same FDR correction procedure was applied for subregion analyses, with correction applied separately within each macroregion-metric combination (i.e., 45 tests per correction, 10 corrections total across 5 macroregions × 2 metrics). Again, results with FDR-adjusted P < 0.05 were considered statistically significant. All data preparation and analyses were conducted using *R* Version 4.5.2 (*78*).

## 3. Results

We tested 45 SNPs (**Tables 1, S1**) previously associated with adult PVS burden or DTI-ALPS diffusion (*32, 33*), as well as AQP4 variants implicated in glymphatic flow and brain-CSF (*34*). These genetic associations were examined against PVS count and volume fraction (VF) across six white matter macroregions and 28 DK atlas subregions (**Table S2**) in 6,600 children aged 9–10 years from the ABCD Study (**Table 2**). Relative to the full ABCD cohort, our sample were from a higher socioeconomic status background, self-reported as White, and were more likely to be scanned on a Siemens MRI scanner via a 64-channel head coil (**Table S3**).

**Table 2.** Sample characteristics. Total number (N) of participants and (percentage of total sample) for each category are presented for the study sample. Mean and standard deviation (SD) are presented for numeric variables. Abbreviations: Adolescent Brain Cognitive Development (ABCD) Study, General Educational Development (GED), General Electric (GE). ^+^Other race/ethnicity category includes participants identified by their caregiver as American Indian/Native American, Alaska Native, Native Hawaiian, Guamanian, Samoan, Other Pacific Islander, Asian Indian, Chinese, Filipino, Japanese, Korean, Vietnamese, Other Asian not listed, or Other Race not listed.

|  | N (% of Sample) |
| --- | --- |
| N | 6,600 |
| Age at baseline (months) | 119.2 (7.5) |
| Sex (% Female) | 3,200 (48.5%) |
| *Race & Ethnicity* |  |
| Non-Hispanic Asian | 47 (0.7%) |
| Hispanic/Latino/a | 1,339 (20.3%) |
| Non-Hispanic Black | 941 (14.3%) |
| Non-Hispanic White | 3,722 (56.4%) |
| Other<sup>+</sup>/Multi-Racial | 550 (8.3%) |
| *Household Income* |  |
| < \$50,000 | 1,647 (25.0%) |
| ≥\$50,000 to <\$100,000 | 1,767 (26.8%) |
| ≥ \$100,000 | 2,641 (40.0%) |
| Don't Know/Refuse to Answer | 545 (8.3%) |
| *MRI Scanner Manufacturer* |  |
| Siemens | 4,474 (67.8%) |
| GE Medical Systems | 1,301 (19.7%) |
| Philips Medical Systems | 825 (12.5%) |
| *Head coil* |  |
| 32-channel | 5,481 (83.0%) |
| 64-channel | 1,119 (17.0%) |

We first characterized the distribution of PVS count and VF across DK subregions to establish the regional landscape against which genetic associations were tested. Both PVS count and VF varied substantially across brain regions (**Figure 1A-B**). The highest PVS counts were observed in the frontal and parietal lobes, with the lowest counts in the occipital and portions of the temporal lobes. VF showed a similar regional pattern. At ages 9-10 years, the occipital lobe had the lowest PVS VF (Mean ± SD = 0.002 ± 0.001), particularly within the lingual and cuneus regions. The frontal lobe had the highest PVS VF (Mean ± SD = 0.006 ± 0.002), specifically within the caudal middle frontal and superior frontal regions. Marked interindividual variability was observed across all regions, quantified by the coefficient of variation (CV), reflecting the degree to which children differ from one another in regional PVS morphology (**Figure 1C-D**). The highest between-subject variability was observed in occipital lobe VF (CV=50%), driven by the cuneus (55%) and lateral occipital (49%) subregions, while the paracentral region showed the highest CV across all subregions (58%; **Table S4, Figure S3**).

**Figure 1.**
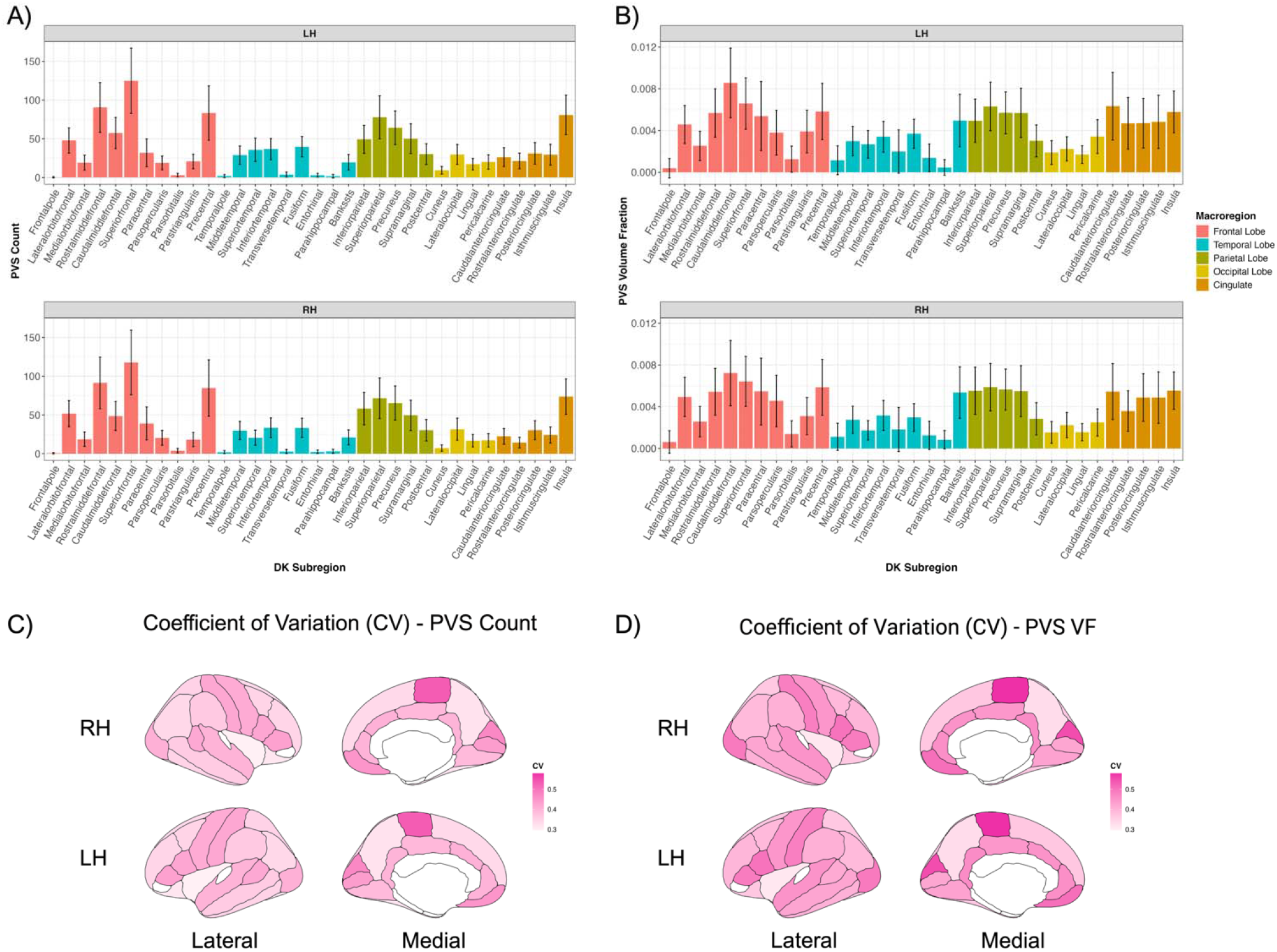
Distributions of perivascular space (PVS) A) Count and B) Volume Fraction (VF) by region and hemisphere. The x-axis depicts each Desikan-Killiany (DK) subregion, color coded by macroregion. Bars reflect mean and error bars reflect standard error. Coefficient of Variation (CV) plots for **PVS C) Count**, and **D) VF**. CV (standard deviation/mean) reflects the degree of between-subject variability in regional PVS morphology relative to the regional mean, allowing comparison of variability across regions with different average PVS levels. Brain maps are rendered such that darker pink reflects higher CV and lighter pink reflects lower CV. Abbreviations: Left Hemisphere (LH), Right Hemisphere (RH), Perivascular Space (PVS). Figure created using R and Biorender.

### 3.1 Genetic Associations with PVS Morphology

Of the 45 SNPs examined, 15 (33%) demonstrated significant associations with PVS morphology in at least one macroregion, with effects expressed as standardized betas to facilitate comparison across PVS outcomes and brain regions (**Figure 2A-B; Tables S5-S6)**. Associations were predominantly observed with PVS VF rather than count measures (43 vs. 4 significant macroregion associations). The four significant count associations represented a subset of findings that also reached significance for VF, but with larger betas in the VF models. Of these 15 FDR-corrected significant SNPs, 11 (46.7%) were intronic variants (**Figure 2C**), and 12 (80%) were identified through the adult PVS burden GWAS (*32*) (**Figure 2D**).

**Figure 2.**
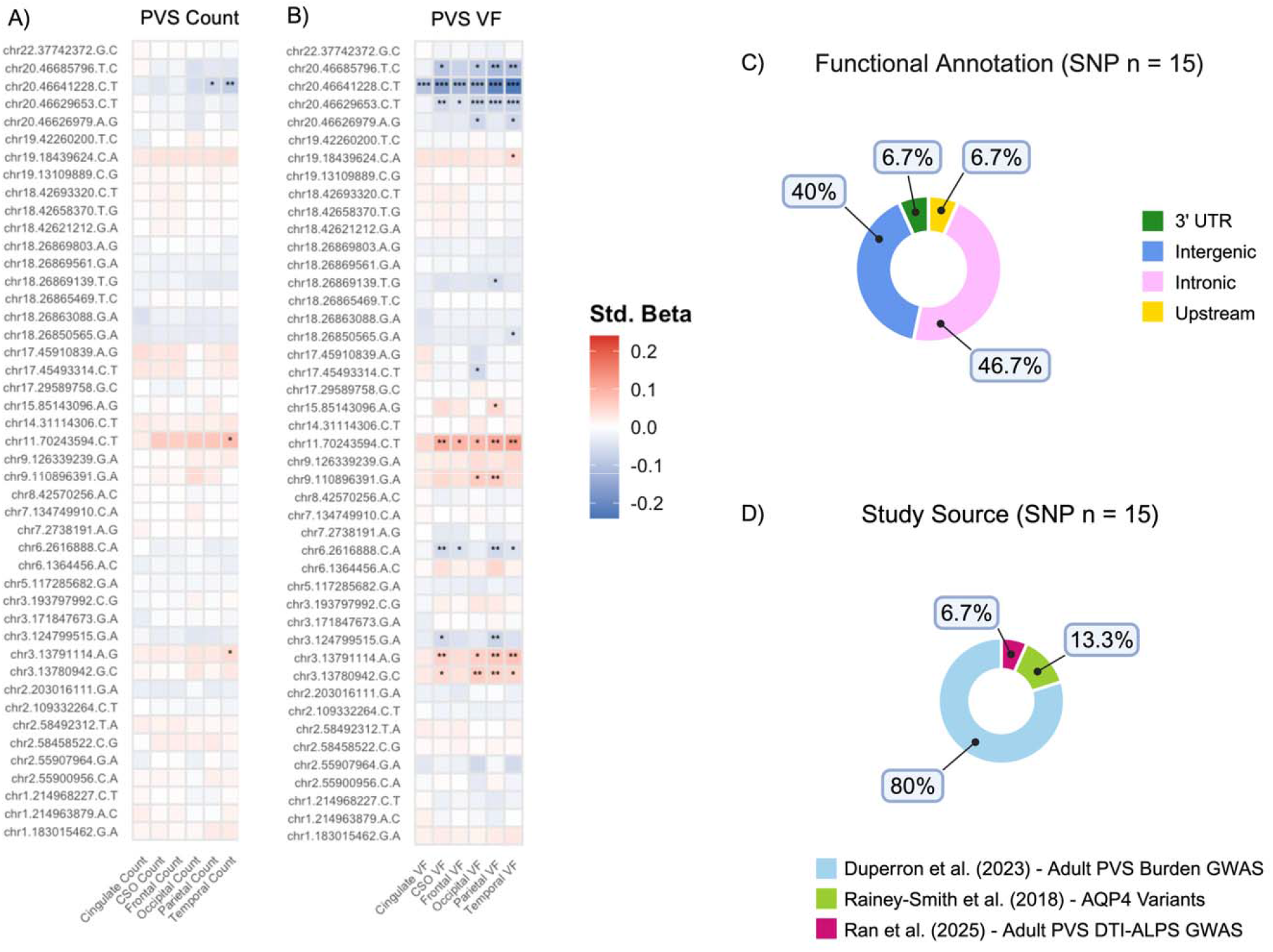
Associations between genetic variants and perivascular space (PVS) **A) count** and **B) volume fraction (VF) in six white matter macroregions**. Macroregions are depicted on each respective x-axis, whereas single nucleotide polymorphisms (SNPs) are listed on the y-axis in chr:pos:ref:alt format in GRCh38 build. SNPs are organized in ascending order from bottom to top (chromosome 1 to 22). Standardized (std.) beta color mapping has been applied to each cell to depict the beta coefficient for each association, and false discovery rate (FDR)-corrected significant cells have been indicated using the following legend: * = P_FDR_ < 0.05, ** = P_FDR_ < 0.01, *** = P_FDR_ < 0.001. Distribution of FDR-significant SNPs by **C) functional annotation**, and **D) study source**. Abbreviations: Centrum Semiovale (CSO), Perivascular Space (PVS), Genome Wide Association Study (GWAS), Diffusion Tensor Im ging Along the Perivascular Space (DTI-ALPS). Figure created using R and Biorender.

The strongest and most widespread effects were observed for SNPs located near the *SLC13A3* gene on chromosome 20. The lead variant, chr20:46641228, was significantly associated with lower PVS VF across all six macroregions (β = −0.069 to −0.23, all P_FDR_ < 0.001), with each additional copy of the minor allele associated with progressively lower PVS VF, such that homozygous carriers showed the lowest PVS VF (**Table S6**). Two additional nearby variants on chromosome 20 demonstrated similarly broad effects on PVS VF: chr20:46629653 showed significant associations across all examined regions (β = −0.047 to −0.078, all P_FDR_ < 0.05), and chr20:46685796 demonstrated associations in the CSO as well as the occipital, parietal, and temporal lobes (β = −0.086 to −0.117, all P_FDR_ < 0.05) (**Table S6**). One chromosome 11 variant (chr11:70243594) near the *CENPF* and *KCNK2* genes demonstrated positive associations across five of the six regions, with the strongest effects in temporal (β = 0.12, P_FDR_ = 0.002) and parietal (β = 0.11, P_FDR_ = 0.004) lobe VF (**Table S6**). Variants on chromosome 3 demonstrated widespread, divergent directional effects. One variant nearby *ITGB5* and *UMPS* genes (chr3:124799515) was associated with lower CSO (β = −0.06, P_FDR_ = 0.02) and parietal (β = −0.07, P_FDR_ = 0.007) VF, while two other variants near the *WNT7A* gene (chr3:13780942 and chr3:13791114) were associated with higher VF in the CSO (β = 0.045 to 0.056, all P_FDR_ < 0.05), occipital (β = 0.058 to 0.062, all P_FDR_ < 0.05), parietal (β = 0.054 to 0.067, all P_FDR_ < 0.01), and temporal (β = 0.051 to 0.073, all P_FDR_ < 0.05) lobe. Several other SNPs showed more selective associations with PVS VF (e.g., chr17:45493314) (**Figure 2B; Table S6**).

### 3.2 Analyses of Regional Variability

Given that PVS morphology varies substantially across white matter subregions, both in the current sample (**Figure 1**) and as previously reported (*27*), and that lobar aggregation may obscure spatially restricted genetic effects, we conducted follow-up subregion-level analyses to characterize the anatomical specificity of genetic associations at finer spatial resolution.

While macroregion analyses identified 15 significant SNP associations with PVS morphology, finer-grained subregional analyses identified 21, including 6 SNPs detectable only at finer spatial resolution (**Figure 3A-B**). Of the 45 SNPs tested, 30 (67%) showed significant regional variability across subregions within a lobe (**Tables S7-S8; Supplemental Methods 1.3**), further demonstrating that coarser spatial aggregation can mask spatially distinct genetic effects (**Figure 3A-B; Tables S9-S13**). SNPs near *SLC13A3* on chromosome 20 again showed the most widespread associations across subregions. Chr20:46641228 demonstrated significant VF associations across 22 DK subregions spanning all lobes examined, with count associations extending across 12 subregions (**Figure 4A)**. Two additional chromosome 20 variants (chr20:46629653 and chr20:46685796) showed similarly widespread subregional patterns, while chr20:46626979 exhibited more spatially restricted associations, primarily within the temporal lobe. Beyond chromosome 20 variants, chr11:70243594 showed the next broadest spatial reach, with significant VF associations across 14 subregions spanning the temporal, parietal, frontal, and cingulate cortex (**Figure 4B**). Variants near *WNT7A* on chromosome 3 also showed broad subregional associations, with chr3:13791114 demonstrating significant positive associations with PVS VF across parietal, temporal and occipital regions (**Figure 4C**). Finally, *AQP4* variants demonstrated broad associations on the locus level; however individual SNPs were more regionally specific (**Figure S4**).

**Figure 3.**
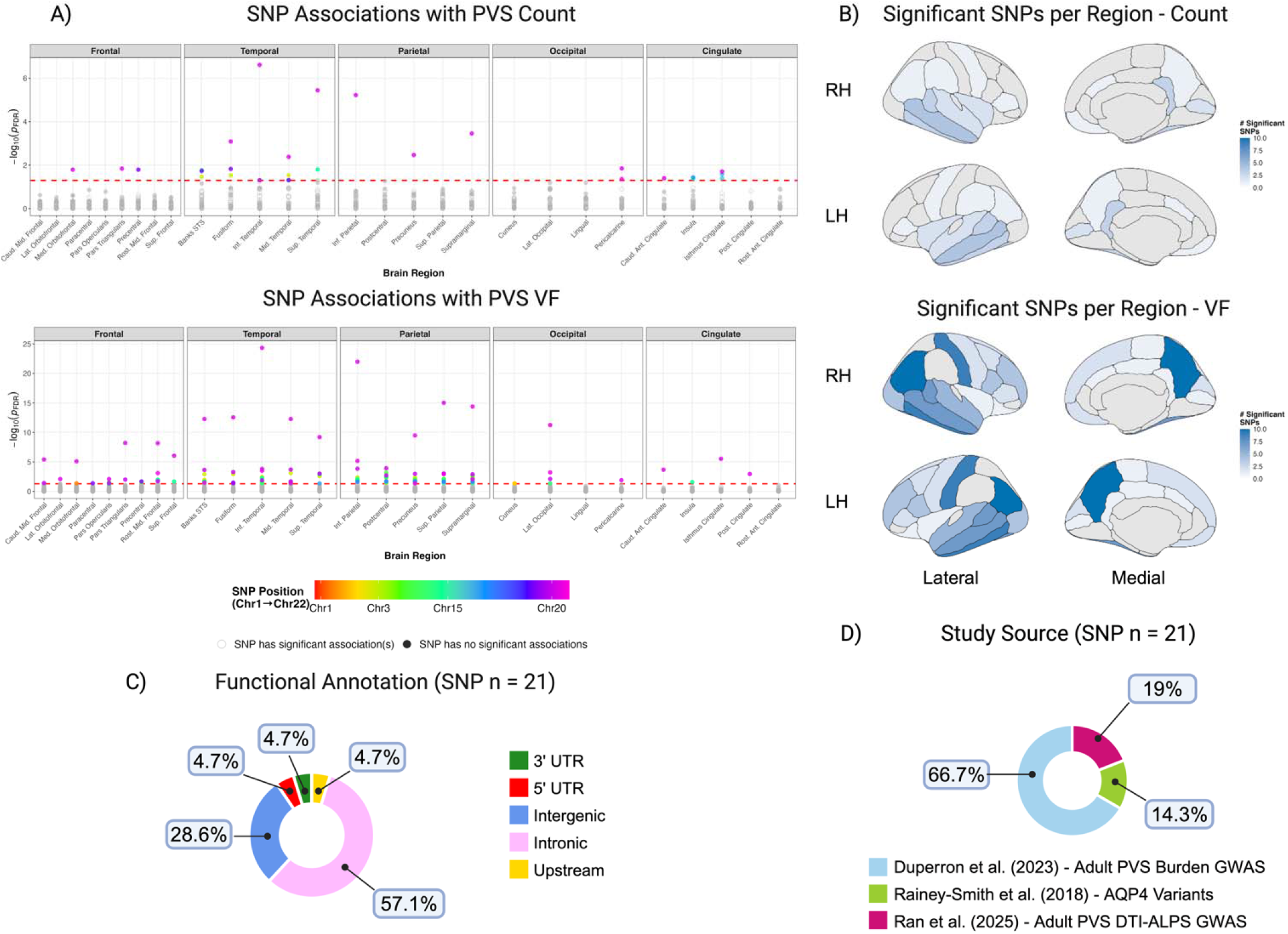
Regional variability in genetic associations with perivascular space (PVS) count and volume fraction (VF). **A)** Each dot reflects an association between one single nucleotide polymorphism (SNP) and each Desikan-Killiany (DK) subregion, with colored dots falling above the red line corresponding to false discovery rate (FDR)-corrected significant associations and grey dots falling below the red line corresponding to null associations. See color bar for legend of SNP color coding. **B) Number of FDR-significant associations per subregion**. Color mapping indicates the number of significant SNPs per DK subregion, where darker blue values represent more significant SNP associations, and lighter blue regions represent fewer significant SNP associations. Grey regions were not significant or were not examined. **Distribution of FDR-significant SNPs by C) functional annotation**, and **D) study source**. Abbreviations: Centrum Semiovale (CSO), False Discovery Rate (FDR), Left Hemisphere (LH), Right Hemisphere (RH), Perivascular Space (PVS), Genome Wide Association Study (GWAS), Diffusion Tensor Imaging Along the Perivascular Space (DTI-ALPS). Figure created using R and Biorender.

**Figure 4.**
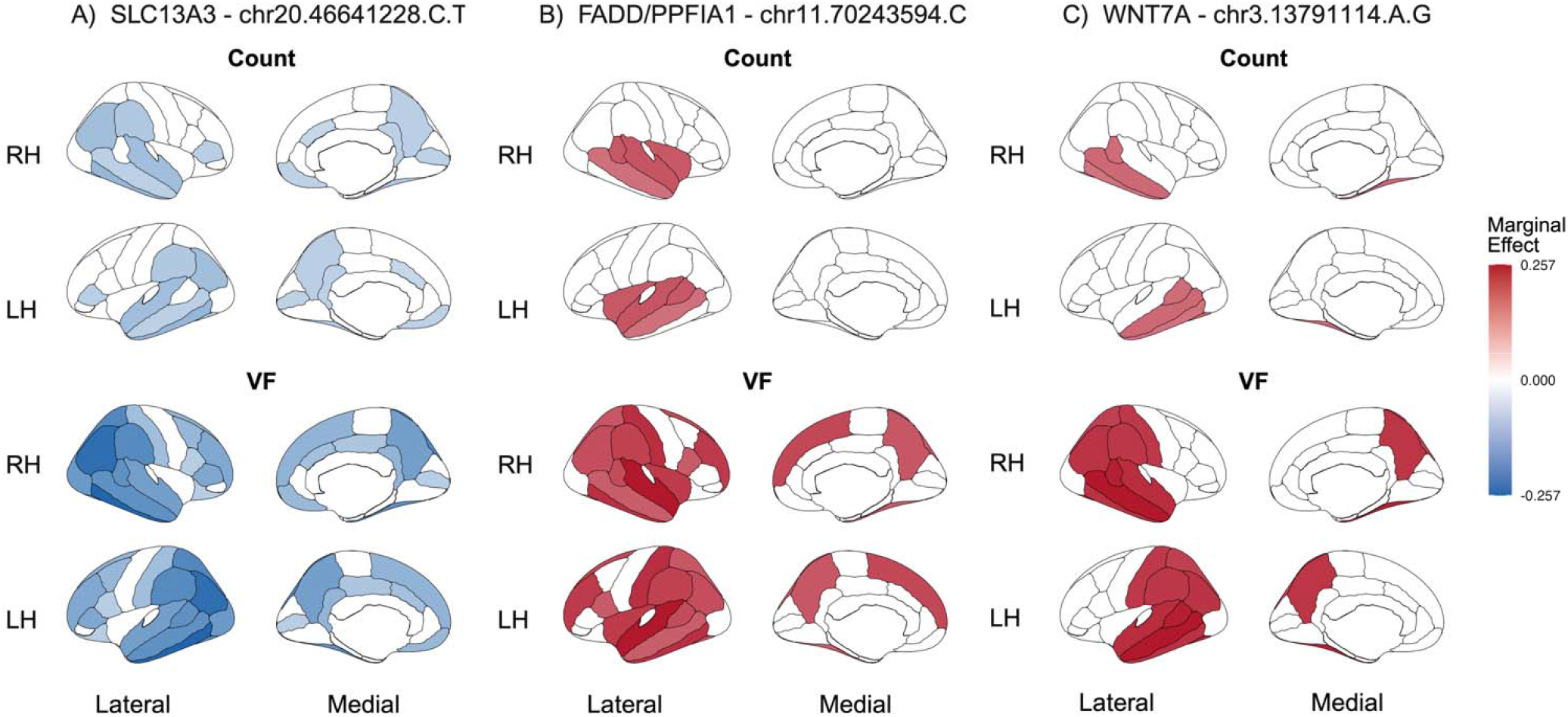
Widespread genetic associations with perivascular space (PVS) subregion morphology. Brain maps depict false discovery rate (FDR)-significant associations between subregion PVS and three single nucleotide polymorphisms (SNPs) showing the broadest spatial reach, including **A) chr20:46641228 (near SLC13A3), B) chr11:70243594 (near *FADD/PPFIA1***), and **C) chr3:13791114 (near *WNT7A*)**. Color mapping reflects the estimated marginal effects for each significant SNP-region associations across Desikan-Killiany (DK) subregions, with darker colors reflecting stronger effects. Red values represent positive associations (i.e., each additional minor allele is associated with a higher PVS count or VF), and blue values represent negative associations (i.e., each additional minor allele is associated with lower PVS count or VF). Abbreviations: Volume Fraction (VF), Left Hemisphere (LH), Right Hemisphere (RH), Perivascular Space (PVS). Figure created using R and Biorender.

Across SNPs, parietal subregions showed the most frequent associations (**Figure 3B**). The supramarginal gyrus, superior parietal cortex, inferior parietal cortex, precuneus, and the postcentral gyrus, each showed significant associations with 9-11 SNPs, suggesting that genetic associations are broadly distributed across the parietal lobe rather than driven by any single subregion (**Figure 3B; Table S10**). Temporal associations were similarly widespread, led by the inferior temporal gyrus (9 SNPs) and the banks STS, fusiform, and middle temporal gyrus (7 SNPs each) (**Figure 3B; Table S9**). In contrast, frontal associations were concentrated in fewer subregions. The pars opercularis and rostral middle frontal gyrus each showed significant associations with 4 SNPs, while most other frontal subregions showed associations driven predominantly by chromosome 20 variants. Occipital and cingulate subregions showed the fewest associations overall, largely confined to the lateral occipital and isthmus cingulate cortex (**Figure 3B**). Of the 21 significant SNPs at the subregion level, 16 (57.1%) were intronic variants (**Figure 3C**), and 14 (66.7%) were identified through the adult PVS burden GWAS (*32*) (*Figure 3D*).

## 4. Discussion

This is the first study to examine genetic associations with PVS morphology in a developing sample, extending adult GWAS findings into a large cohort of pre-adolescents prior to the accumulation of vascular risk factors and age-related neurological change. Nearly half of the tested SNPs replicated in our sample, the majority (14/21, 67%) derived from the adult PVS burden GWAS (*32*). Genetic effects were stronger and more widespread for PVS VF than count at ages 9-10 years. Replicated SNPs converged on a small number of biological pathways whose effects were anatomically patterned, revealing both developmental conservation of specific genetic mechanisms and substantial regional specificity in how those mechanisms may operate.

Replicated SNPs converged on several biological pathways relevant to glymphatic architecture, including solute transport, vascular integrity, and programmed cell death. The strongest and most widespread associations were obs rved for variants near *SLC13A3* on chromosome 20, which encodes a sodium-dependent dicarboxylate cotransporter expressed in the kidneys, choroid plexus, and astrocytes, structures central to CSF production and waste clearance (*39*). *SLC13A3* variants have previously been linked to acute reversible leukoencephalopathy with increased urinary alpha ketoglutarate, a condition characterized by temporary changes to white matter and neurological symptoms (*40*). The negative associations observed here suggest that *SLC13A3* variation may be linked to lower PVS volume, potentially impairing glymphatic solute transport, though the precise mechanism will require functional studies to establish. Critically, *SLC13A3* was originally identified through a GWAS of PVS burden in adults (*32*), yet the detectability of these associations at ages 9–10 indicates that this genetic influence shapes brain clearance architecture during development itself, not merely as a downstream consequence of aging or vascular risk factor accumulation.

A chromosome 11 intergenic variant (chr11:70243594) showed a positive association with PVS VF across nearly the entire brain. Located between *FADD* and *PPFIA1*, this variant may exert regulatory effects on one or both genes through noncoding DNA elements that control gene expression (*41*). *FADD* encodes a critical adaptor protein involved in programmed cell death and immune signaling, while *PPFIA1* encodes a scaffolding protein essential for cell-matrix interactions, synapse organization, and focal adhesion dynamics (*42*). Either pathway could plausibly influence perivascular structure, such as *FADD* through neuroinflammatory regulation of vascular remodeling or *PPFIA1* through its role in maintaining the structural integrity of the neurovascular unit.

Three SNPs on chromosome 3, previously identified in association with PVS burden in adults (*32*), were also associated with PVS VF in the present sample. Two intergenic *WNT7A* variants (chr3:13780942, chr3:13791114) exhibited widespread positive associations, such that individuals carrying more copies of the minor allele demonstrated higher PVS VF. *WNT7A* encodes Wnt7a, a secreted signaling molecule with a well-established role in embryonic development, particularly in limb patterning and neural stem cell proliferation (*43*). Wnt7a activates the Wnt/β-catenin signaling cascade, which plays a fundamental role in BBB formation, angiogenesis, and vascular development (*44*), making *WNT7A* a compelling candidate for genetic analyses of PVS architecture. In contrast, the third chromosome 3 SNP (chr3:124799515), an intronic *ITGB5* variant, showed more localized negative associations. *ITGB5* encodes an integrin subunit involved in cell-extracellular matrix adhesion, tumor progression, and angiogenesis, with particularly high expression in glioblastoma (*45*). Its localized negative associations suggest a potentially more regionally restricted role in perivascular structural integrity. The contrasting directionality and regional specificity of these chromosome 3 associations suggest that even within a single chromosomal region, distinct loci may influence PVS morphology through divergent biological mechanisms. Together, replicated SNPs implicate a core set of processes, including glymphatic solute transport, BBB maintenance, vascular signaling, and cell-matrix adhesion, that appear to be active determinants of PVS morphology during development. The detectability of these associations at ages 9–10, prior to the vascular risk factor accumulation that characterizes adult cohorts, suggests they reflect fundamental biological variation in glymphatic architecture rather than disease-related secondary change.

SNP associations with PVS were not uniformly distributed across subregions within a lobe, and this regional heterogeneity was most pronounced in the frontal lobe. The pronounced frontal heterogeneity likely reflects the anatomical and cellular complexity of the neurovascular unit (NVU) across this lobe (*46*). The frontal lobe contains highly diverse vascular territories with distinct patterns of penetrating vessels, variable capillary density, and region-specific endothelial molecular signatures (*46, 47*), all of which may modulate how genetic variants influence local perivascular structure. Beyond endothelial heterogeneity, other NVU components exhibit similarly pronounced regional variation in order for regional modifications in blood flow to match the energy demand of neural activity (*47*). Astrocytes, which are critical for BBB function (*48*), demonstrate regional heterogeneity in gene expression. For example, mouse studies have demonstrated that astrocytes in several neuroanatomical structures and white matter regions exhibit distinct transcriptional and molecular profiles (*49–51*), and that this region-specific astrocyte gene expression changes across development (*52*). Pericytes, which regulate cerebral blood flow and maintain BBB integrity, show regional differences in density, morphology, and molecular identity (*47, 53*). Microglia also exhibit regional variability, with frontal microglia differing substantially from those in subcortical structures like the thalamus (*54*) Together, the regional diversity of NVU cellular composition provides a plausible biological basis for the anatomically patterned genetic effects observed across white matter subregions.

Regional differences in metabolic demand provide another potential explanation for the heterogeneity in genetic associations across the brain. The prefrontal cortex, for instance, has the highest density of glial cells relative to neurons in the brain, reflecting elevated metabolic demands (*55*). These elevated demands are supported in part by aerobic glycolysis, which accounts for 30% of glucose consumption at birth but declines to approximately 10% in adulthood (*56, 57*). This decline is not uniform across development or across regions; the metabolic transition from aerobic glycolysis to oxidative phosphorylation completes earlier in primary sensory and motor cortices and considerably later in higher-order association cortices, particularly in the prefrontal cortex (*56*). Critically, at ages 9-10, children are in the declining phase of peak glucose uptake, but have not yet reached adult metabolic profiles, meaning that association cortices and primary cortices are likely at different stages of this metabolic transition. Aerobic glycolysis actively drives the biosynthesis of lipids, proteins, and nucleotides needed for neural and vascular membrane maintenance and remodeling, such that regions with persistently elevated aerobic glycolysis impose a higher and more prolonged angiogenic demand on the local vasculature (*56*). These regional differences in metabolic demand drive differences in vascular architecture (*58, 59*) and neurovascular cell composition (*47*), providing a plausible biological basis for the anatomically patterned genetic associations with PVS morphology observed here.

Regional variation in protein expression provides yet another potential mechanism underlying the anatomically patterned genetic associations observed here. For instance, AQP4, a water channel protein critical for glymphatic clearance, shows a range of expression levels across the cortex (∼170-430 normalized transcripts per million [nTPM]), with particularly pronounced heterogeneity within the frontal lobe (*60*), which may help explain why *AQP4* SNP associations with frontal PVS were sparse and subregion-specific rather than uniform. Recent work further shows that the relationship between AQP4 protein levels and Aβ, a toxic protein cleared by the glymphatic system and implicated in Alzheimer’s disease, varies by brain region (*61*), supporting the notion that genetic determinants of the glymphatic system may be regionally specific. Collectively, this heterogeneous neurovascular landscape, encompassing variability in capillary density, glucose metabolism, endothelial molecular signatures, and AQP4 expression, provides a plausible biological basis for the anatomically patterned genetic effects observed here, such that only the most spatially widespread SNPs produced consistent associations across regions.

Subregion-level analyses revealed complementary patterns of regional specificity. Despite showing the greatest variability in SNP effect magnitudes across subregions, the frontal lobe was associated with relatively few SNPs overall. Instead, parietal and temporal subregions showed the most widespread associations, with every subregion within these lobes implicated by at least one SNP for VF. Parietal subregions showed the most consistent genetic associations across all SNPs: the supramarginal gyrus, superior and inferior parietal cortex, and precuneus each showed associations with 9–11 SNPs. Temporal associations were similarly distributed across subregions, while frontal, occipital, and cingulate associations were each anchored by a small number of specific subregions.

Together, these findings suggest that the regional landscape of genetic effects on PVS in childhood likely reflects an interplay between local neurovascular complexity and developmental timing. The frontal lobe’s heterogeneous vascular environment may restrict detectable associations to genetic variants affecting biological processes shared broadly across its diverse subregions. Conversely, the uniformly high metabolic demands of actively maturing parietal and temporal cortex, undergoing rapid myelination and cortical thinning simultaneously across subregions (*62, 63*), may create a more receptive biological context in which a broader range of genetic variants can exert detectable effects. The frontal lobe is also the last cortical region to mature and, at ages 9-10, may not yet have reached the developmental stage at which these genetic influences are most strongly expressed, in contrast to the parietal and temporal cortices, which are undergoing their most active period of change at precisely this age. Whether frontal genetic effects emerge more strongly later in development, as the frontal lobe reaches its own maturational peak, and whether parietal and temporal associations persist beyond this active window, remain important questions for longitudinal and cross-age studies to address. Critically, this regional specificity was detectable only through fine-grained subregion analyses, signals that would have been missed at the lobe or whole brain level, underscoring that the spatial resolution of imaging analyses is not merely a methodological choice but a biological one.

Lastly, the pattern of replication across source studies is itself informative. Nearly all replicated SNPs were derived from the adult PVS burden GWAS which used structural MRI (*32*), while few SNPs replicated from the DTI-ALPS GWAS (*33*). This may reflect a genuine biological distinction between phenotypes. The adult PVS burden GWAS (*32*) quantified PVS using visual semiquantitative rating scales dichotomized into extensive versus non-extensive burden, a coarser measure that captures extreme PVS load rather than continuous morphological variation. The DTI-ALPS GWAS (*33*) measured diffusion tensor imaging along the perivascular space to capture interstitial fluid movement rather than PVS structure directly (*38*). In contrast, the present study used an automated segmentation pipeline to quantify continuous PVS count and volume fraction across 28 subregions, capturing fine-grained regional morphological variation at a spatial resolution unavailable in either source study. That replication was highest with the adult PVS burden GWAS (*32*), despite its coarser phenotyping, may reflect shared genetic determinants of PVS morphology across the lifespan, while the lower replication with DTI-ALPS (*33*) may indicate that diffusivity along the PVS and structural PVS morphology, are not genetically similar phenotypes. These distinctions underscore the importance of phenotype definition in genetic studies of the glymphatic system, as different measures may capture distinct biological dimensions with partially non-overlapping genetic architectures. Future work examining both structural measures and fluid dynamics of PVS within the same pediatric sample are needed to clarify whether the genetic architectures underlying PVS morphology and function are shared, and at what developmental stage they may diverge.

### 4.1 Strengths & Limitations

This study has several strengths. The use of a large, diverse cohort with harmonized MRI and genetic data provided sufficient power to detect associations across multiple regions and metrics, while the cross-ancestry sample and inclusion of genetic ancestry principal components enhanced the generalizability of findings beyond European-ancestry populations.

Several limitations should also be considered. First, this is a candidate SNP study rather than a GWAS, as the ABCD Study does not have the required sample size for a well powered GWAS. While testing 45 SNPs provides valuable replication and extension of previous findings, it offers a narrow window into the full genetic architecture of PVS morphology, and novel SNPs important specifically during adolescence may have been missed. Second, nearly all tested SNPs are located in non-coding regions of the genome (i.e., intronic, intergenic, or UTR variants). Non-coding variants can exert regulatory effects on gene expression, but identifying causal genes and uncovering the functional consequences of these variants will require expression quantitative trait locus analyses and experimental validation. Third, six brain regions (the frontal pole, pars orbitalis, parahippocampal, entorhinal, transverse temporal, and temporal pole) were excluded due to insufficient PVS signal for reliable quantification at the current MRI resolution, meaning that genetic associations in these regions remain uncharacterized and represent an important target for future work as imaging methods improve. Finally, as a cross-sectional study, we cannot determine whether the genetic associations identified here are stable across development or specific to this window of rapid neural change, a question that only longitudinal designs can address.

## 5. Conclusion

This study provides the first evidence that genetic influences on PVS morphology are detectable in childhood and that these effects are, at least in part, developmentally conserved. By demonstrating that nearly 50% of adult-identified SNPs, particularly those near *SLC13A3, WNT7A, FADD/PPFIA1*, and *AQP4*, associate with PVS VF at ages 9–10, we show that the genetic architecture of PVS morphology is established in childhood, before the onset of age-related vascular pathology. Equally important, the regional specificity of these associations demonstrates that genetic effects on PVS are anatomically patterned and cannot be fully captured through coarse regional analyses alone. The frontal lobe is characterized by a few SNPs whose effect magnitudes vary substantially across subregions, whereas parietal and temporal cortex show broader genetic involvement across many SNPs and subregions, a distinction that only becomes apparent at fine spatial resolution.

## Acknowledgments

A special thank you to all participants and their families for their participation in the ABCD Study. Data used in the preparation of this article were obtained from the Adolescent Brain Cognitive Development^SM^ (ABCD) Study (https://abcdstudy.org), held in the NIMH Data Archive (NDA). This is a multisite, longitudinal study designed to recruit more than 10,000 children aged 9-10 and follow them over 10 years into early adulthood. The ABCD Study® is supported by the National Institutes of Health and additional federal partners under award numbers U01DA041048, U01DA050989, U01DA051016, U01DA041022, U01DA051018, U01DA051037, U01DA050987, U01DA041174, U01DA041106, U01DA041117, U01DA041028, U01DA041134, U01DA050988, U01DA051039, U01DA041156, U01DA041025, U01DA041120, U01DA051038, U01DA041148, U01DA041093, U01DA041089, U24DA041123, U24DA041147. A full list of supporters is available at https://abcdstudy.org/federal-partners.html. A listing of participating sites and a complete listing of the study investigators can be found at https://abcdstudy.org/consortium_members/. ABCD consortium investigators designed and implemented the study and/or provided data but did not necessarily participate in the analysis or writing of this report. This manuscript reflects the views of the authors and may not reflect the opinions or views of the NIH or ABCD consortium investigators. The ABCD data repository grows and changes over time. The ABCD data used in this report came from DOI: 10.15154/8873-zj65 and DOI: 10.15154/1520591.

## Funding

This work was supported by the National Institutes of Health (NIH) National Institute of Environmental Health Sciences (NIEHS) (Grant Nos. P30ES07048 [to JC/MMH]; T32ES013678 [to JM]; P30ES07048 [to WG]); National Institute of Mental Health (NIMH) (Grant RF1MH123223 [to JC]), and National Institute of Neurological Disorders and Stroke (Grant R01NS128486 [to JC]), as well as the USC Office of Research and Innovation – Research Catalyst Program.

## Author Contributions

Conceptualization: JM, MMH

Methodology: JM, HA, JC

Formal analysis: JM

Resources: MMH, JC

Data curation: JM, CT, RC, HL

Writing - original draft: JM

Writing - review and editing: JM, MMH, JC, CT, HA, RC, JG, HL

Visualization: JM

Supervision: MMH, JC

Project administration: MMH

Funding acquisition: JM, MMH, JC

## Competing Interests

The authors declare no competing interests.

## Data and Materials Availability

Data used in the preparation of this article were obtained from the Adolescent Brain Cognitive Development (ABCD) Study (https://abcdstudy.org), held in the NIMH Data Archive (NDA), releases 3.0 (DOI: 10.15154/8873-zj65) and 5.0 (DOI: 10.15154/1520591). Individual participant data will not be made available. Code will be made available upon request.

## Notes

https://osf.io/2yzwe/overview

### Competing Interest Statement

The authors have declared no competing interest.

