## Supplement for "Genetic Architecture of Perivascular Space Morphology in the Pediatric Brain"

**1. Supplemental Methods**

**1.1 Macroregion Analyses**

We conducted linear mixed-effects (LME) models using the *lme4* package (*1*) in *R* Version 4.5.2. These models tested the association between each single nucleotide polymorphism (SNP; N = 45) dosage (continuous 0 to 2 scale) and each perivascular space (PVS) outcome (N = 12). Outcome variables were standardized prior to running each model to aid in interpretability of β coefficients. Models were of the following form per each of the six macroregions (i.e., frontal lobe, temporal lobe, parietal lobe, occipital lobe, the cingulate, and centrum semiovale (CSO)):

*PVS Count ~ 1 + SNP Dosage + Age + Sex + Household Income + Scanner/Coil + Ancestry PCs (1-5) + Regional Volume + (1 | Site)*

*PVS Volume Fraction ~ 1 + SNP Dosage + Age + Sex + Household Income + Scanner/Coil + Ancestry PCs (1-5) + (1 | Site)*

Cohen’s f^2^ partial effect sizes were calculated using the *effectsize* package (*2*) to determine the proportion of variance in each outcome explained by the SNP.

**1.2 Model Diagnostics**

Model assumptions were evaluated for both macroregion and subregion analyses using the *performance* package (*3*) and through visual inspection of diagnostic plots, including but not limited to linearity (residuals vs. fitted plots) and residual normality (Q-Q plots) for each level of residuals. In macroregion analyses, one outcome, occipital VF, demonstrated a slight right skew leading to heavy tails in its level one residual plot. Consequently, we log-transformed this outcome to satisfy linear regression assumptions. All other outcomes met assumptions without any data manipulation.

In subregion analyses, six Desikan-Killiany (DK) regions (the frontal pole, pars orbitalis, parahippocampal gyrus, entorhinal cortex, transverse temporal cortex, and temporal pole) exhibited excessive zero values suggesting a data censoring issue, likely reflecting technical limitations in PVS detection in these regions rather than true biological patterns. Considering that including these regions would introduce significant measurement error and statistical modeling challenges, we excluded these regions from our post-hoc analyses and instead focused on the remaining 28 bilateral cortical regions with reliable PVS quantification. Model diagnostics for the subregion analyses indicated that residual heteroskedasticity and nonlinearity were driven by between-region variability. Therefore, we implemented within-region standardization prior to analyses to meet regression assumptions.

**1.3 Subregional Analyses**

Subregion analyses included 28 bilateral DK regions grouped into five of the macroregions. All outcomes were standardized prior to analysis to enable comparison across subregions. Models took the following form:

*PVS Count ~ SNP + SNP:DK_Subregion_ + Age + Sex + Household Income + Scanner/Coil + Ancestry PCs (1-5) + Regional Volume + (1 | Site) + (1 | Subject)*

*PVS Volume Fraction ~ SNP + SNP:DK_Subregion_ + Age + Sex + Household Income + Scanner/Coil + Ancestry PCs (1-5) + (1 | Site) + (1 | Subject)*

Region main effects were excluded because outcome standardization centered each region at zero, absorbing region-level mean variation and rendering region intercepts uninterpretable. FDR correction was applied separately within each macroregion and metric combination (45 tests per correction), preserving power to detect region-specific effects while controlling for multiple comparisons. Interactions surviving FDR < 0.05 were considered significant. Significant SNP × subregion interactions indicate that genetic effects differ in magnitude and/or direction across specific DK regions within a macroregion, suggesting region-specific genetic architecture of PVS burden. Thus, estimated marginal trends (emmeans::emtrends()) (*4*) were used to estimate the SNP effect within each DK subregion separately, holding covariates at their reference values. These subregion-specific effects were again FDR-corrected within each macroregion and metric combination, identifying which subregions drove the interaction and quantifying the magnitude of genetic associations within each region independently.

**Tables (in a separate excel spreadsheet)**

Table S1. Gene and function mapping for single nucleotide polymorphisms (SNPs) previously associated with perivascular space (PVS) structure, diffusion, or glymphatic function in adults. SNPs presented along with their functional annotation (i.e., most severe consequence) and nearest gene, derived from either the source paper or the Ensembl Variant Predictor tool, along with each gene’s function (from the National Center for Biotechnology Information (NCBI) database), and the brain outcome each SNP was previously associated with in adults. Abbreviations: White Matter (WM), Aquaporin-4 (AQP4), Basal Ganglia (BG), Blood Brain Barrier (BBB).

Table S2. Regions of Interest. Abbreviations: White Matter (WM), Left Hemisphere (LH), Right Hemisphere (RH). Macroregion volume or count (column A) calculated by adding the volume or count of PVS in the following Desikan-Killiany (DK) regions (column B).

Table S3. Comparisons between whole ABCD cohort and current study sample. Differences in sample characteristics between the study sample and whole ABCD sample have been tested using Pearson’s Chi-Square or t-tests, as appropriate. Stars in column A indicate test significance, where * = p < 0.05, ** = p < 0.01, and *** = p < 0.001. Items without stars (i.e., Sex and Age) were not statistically significantly different between samples.

Table S4. Descriptive statistics for perivascular space (PVS) outcomes. Mean, standard deviation (SD), and coefficient of variation (CV) calculated for PVS count and volume fraction (VF) in each of the six macroregions examined. CV calculated by dividing the SD/Mean and multiplying by 100 to obtain a percentage of variation, allowing for an estimation of relative dispersion, or variability.

Table S5. Linear Mixed-Effects modeling results for perivascular space (PVS) count at the macroregion level. Rows highlighted yellow indicate false discovery rate (FDR) significant single nucleotide polymorphism (SNP) - brain associations. Abbreviations: Standardized (Std.), Confidence Interval (CI).

Table S6. Linear Mixed-Effects modeling results for perivascular space (PVS) volume fraction (VF) at the macroregion level. Rows highlighted yellow indicate false discovery rate (FDR) significant single nucleotide polymorphism (SNP) - brain associations. Abbreviations: Standardized (Std.), Confidence Interval (CI).

Table S7. ANOVA results from post-hoc linear mixed-effects analyses of SNP-by-DK region interactions on perivascular space (PVS) count within each of the six macrogregions of interest. Rows highlighted yellow indicate false discovery rate (FDR) significant single nucleotide polymorphism (SNP) - brain associations. Abbreviations: Standardized (Std.), Confidence Interval (CI).

Table S8. ANOVA results from post-hoc linear mixed-effects analyses of SNP-by-DK region interactions on perivascular space (PVS) volume fraction (VF) within each of the six macrogregions of interest. Rows highlighted yellow indicate false discovery rate (FDR) significant single nucleotide polymorphism (SNP) - brain associations. Abbreviations: Standardized (Std.), Confidence Interval (CI).

Table S9. Frontal lobe: Post-hoc SNP associations with perivascular space (PVS) morphology across Desikan-Killiany (DK) frontal lobe subregions, derived from estimated marginal trends of significant SNP-by-subregion interactions in linear mixed-effects models. Rows highlighted yellow indicate false discovery rate (FDR) significant single nucleotide polymorphism (SNP) - brain associations. Abbreviations: Standardized (Std.), Lower Confidence Limit (LCL), Upper Confidence Limit (UCL).

Table S10. Temporal lobe: Post-hoc SNP associations with perivascular space (PVS) morphology across Desikan-Killiany (DK) temporal lobe subregions, derived from estimated marginal trends of significant SNP-by-subregion interactions in linear mixed-effects models. Rows highlighted yellow indicate false discovery rate (FDR) significant single nucleotide polymorphism (SNP) - brain associations. Abbreviations: Standardized (Std.), Lower Confidence Limit (LCL), Upper Confidence Limit (UCL).

Table S11. Parietal lobe: Post-hoc SNP associations with perivascular space (PVS) morphology across Desikan-Killiany (DK) parietal lobe subregions, derived from estimated marginal trends of significant SNP-by-subregion interactions in linear mixed-effects models. Rows highlighted yellow indicate false discovery rate (FDR) significant single nucleotide polymorphism (SNP) - brain associations. Abbreviations: Standardized (Std.), Lower Confidence Limit (LCL), Upper Confidence Limit (UCL).

Table S12. Occipital lobe: Post-hoc SNP associations with perivascular space (PVS) morphology across Desikan-Killiany (DK) occipital lobe subregions, derived from estimated marginal trends of significant SNP-by-subregion interactions in linear mixed-effects models. Rows highlighted yellow indicate false discovery rate (FDR) significant single nucleotide polymorphism (SNP) - brain associations. Abbreviations: Standardized (Std.), Lower Confidence Limit (LCL), Upper Confidence Limit (UCL).

Table S13. Cingulate and Insula: Post-hoc SNP associations with perivascular space (PVS) morphology across Desikan-Killiany (DK) cingulate and insula subregions, derived from estimated marginal trends of significant SNP-by-subregion interactions in linear mixed-effects models. Rows highlighted yellow indicate false discovery rate (FDR) significant single nucleotide polymorphism (SNP) - brain associations. Abbreviations: Standardized (Std.), Lower Confidence Limit (LCL), Upper Confidence Limit (UCL).

**Figures**

**Figure S1. Flowchart of participant exclusion**. Abbreviations: Adolescent Brain Cognitive Development Study (ABCD), Magnetic Resonance Imaging (MRI), Quality Control (QC), Perivascular Space (PVS), Enhanced PVS Contrast image (EPC).

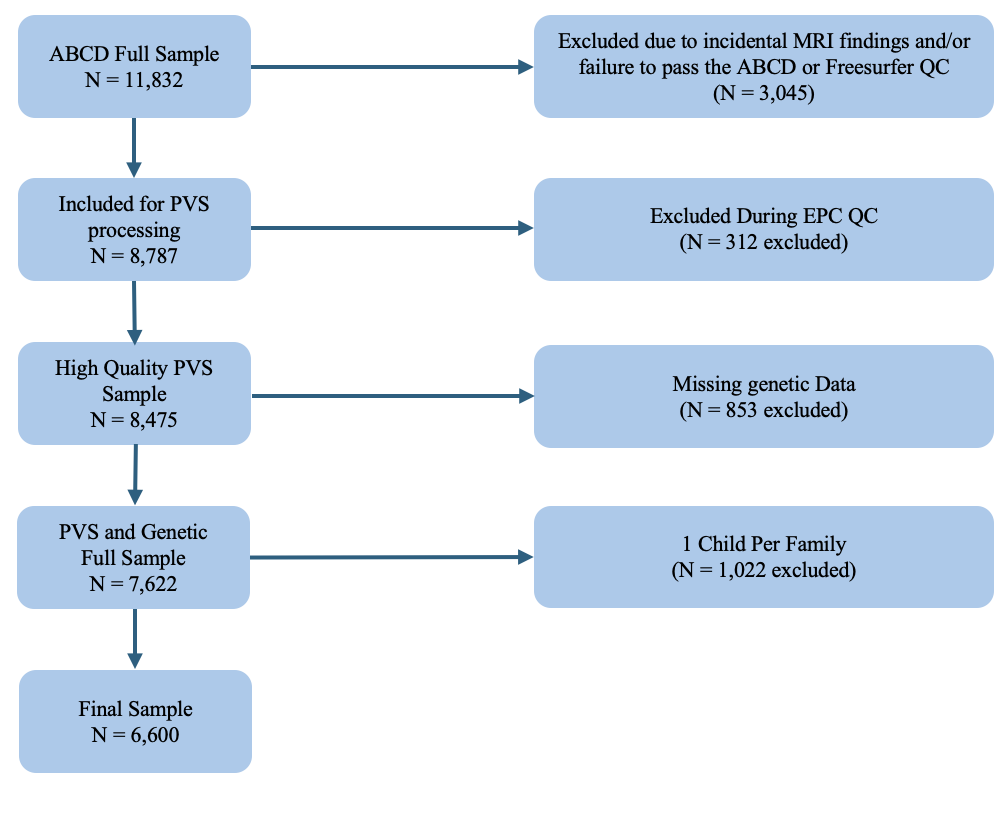

**Figure S2. Pairwise scatterplots of the first five ancestry principal components (PCs) colored by self-reported race/ethnicity.** Each panel displays a pairwise comparison of ancestry PCs 1-5 derived for the ABCD Sample using PC-AiR from the GENESIS package. Points are colored by self-reported race/ethnicity. Clustering across PC pairs indicates that the top 5 PCs effectively capture the major axes of genetic ancestry variation within the sample.

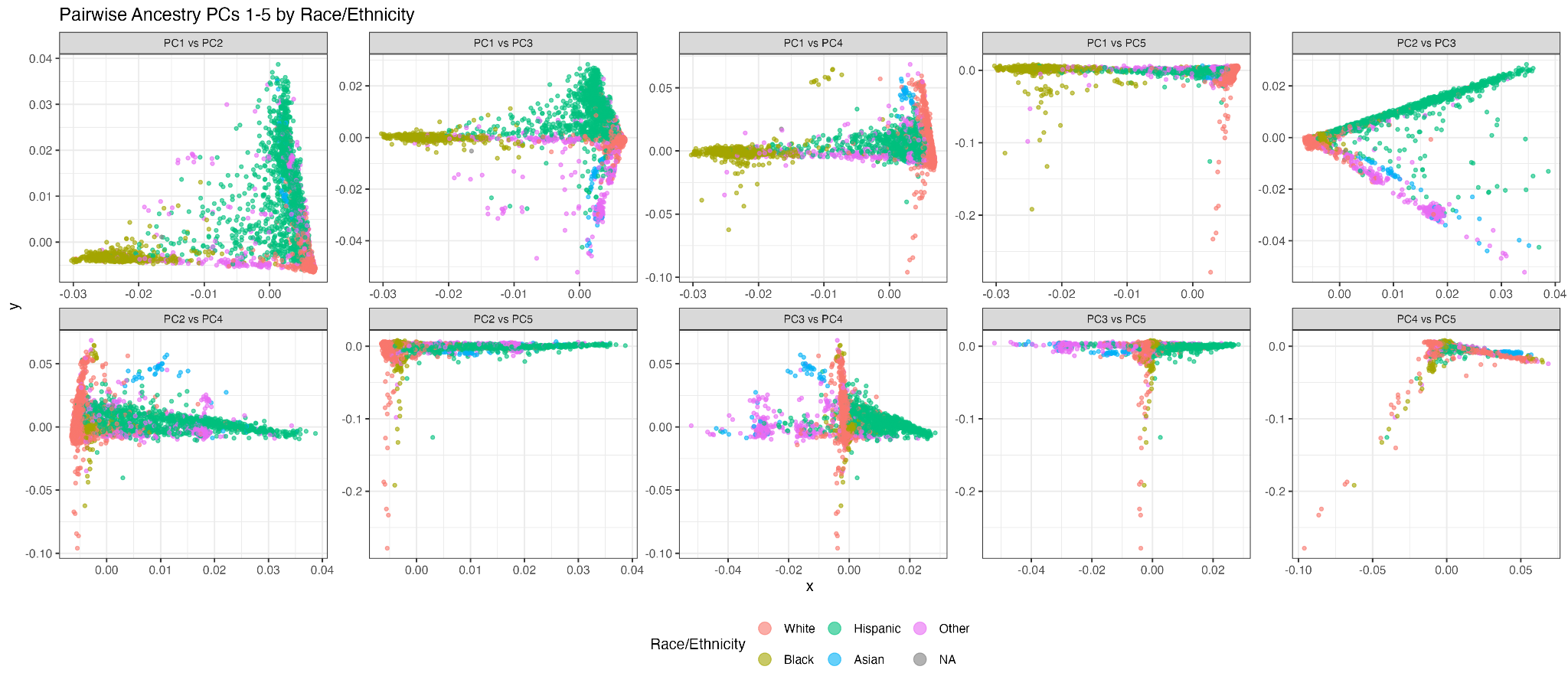

**Figure S3. Perivascular space (PVS) volume fraction (VF) distributions across Desikan-Killiany (DK) atlas regions by lobe.** Ridge plots display the distribution of PVS VF for each cortical region defined by the DK atlas, organized by lobe. For each region, PVS VF values were summed across left and right hemispheres to yield a bilateral measure. Each ridge represents the kernel density estimate of bilateral PVS VF across participants. Regions are ordered by lobe along the y-axis.

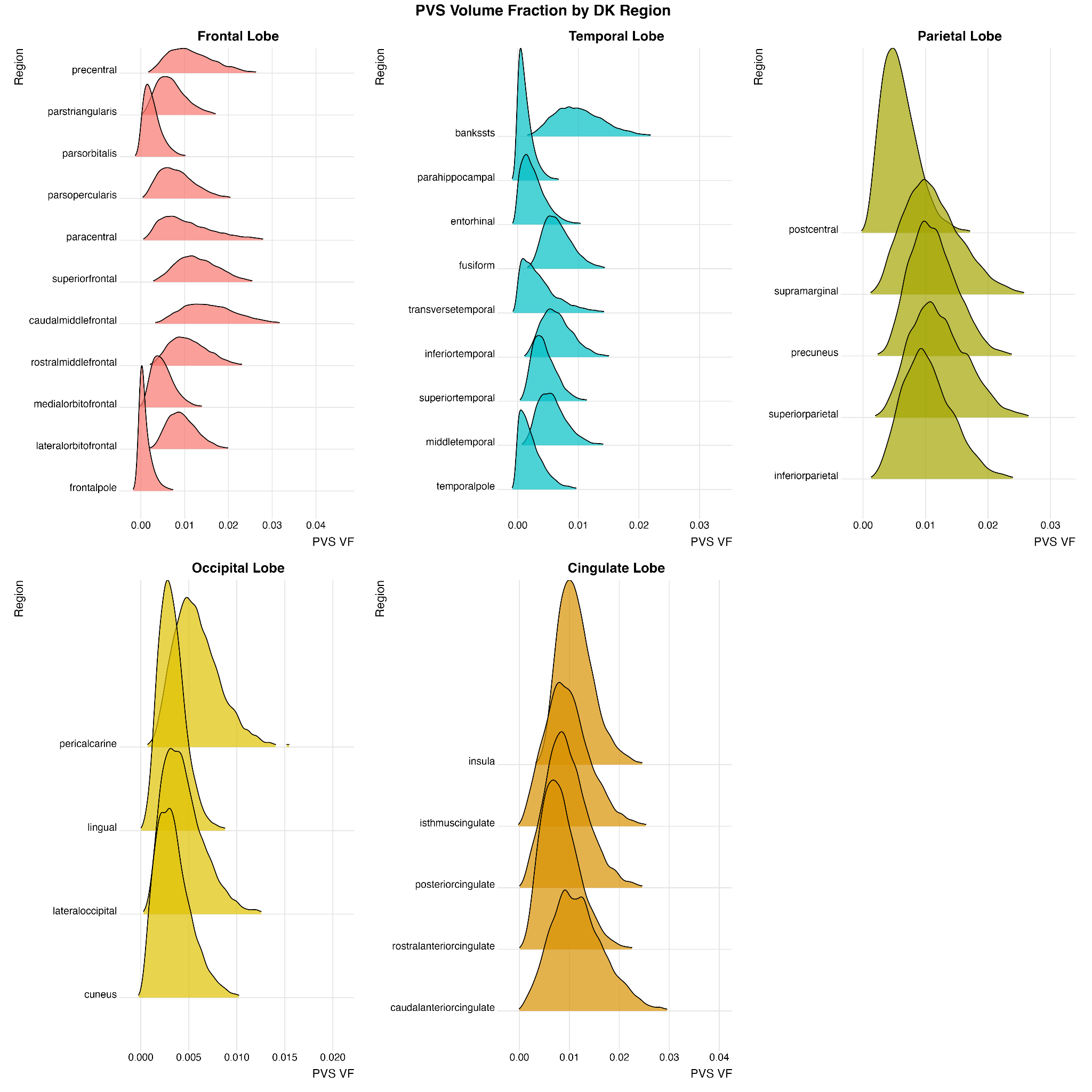

**Figure S4. False discovery rate (FDR)-significant associations between aquaporin-4 (AQP4) single nucleotide polymorphisms (SNPs) and subregion-level perivascular space (PVS) count and volume fraction (VF).** Color mapping reflects where each AQP4 SNP displays an FDR-significant association across Desikan-Killiany subregions. The green region represents the significant association between chr18:26863088 and insula count; the red region represents the significant association between chr18:26850565 and superior temporal VF; the blue regions represent significant associations between chr18.26869139 and parietal VF (inferior parietal, postcentral, precuneus, superior parietal, and supramarginal). White regions are non-significant or were not examined.

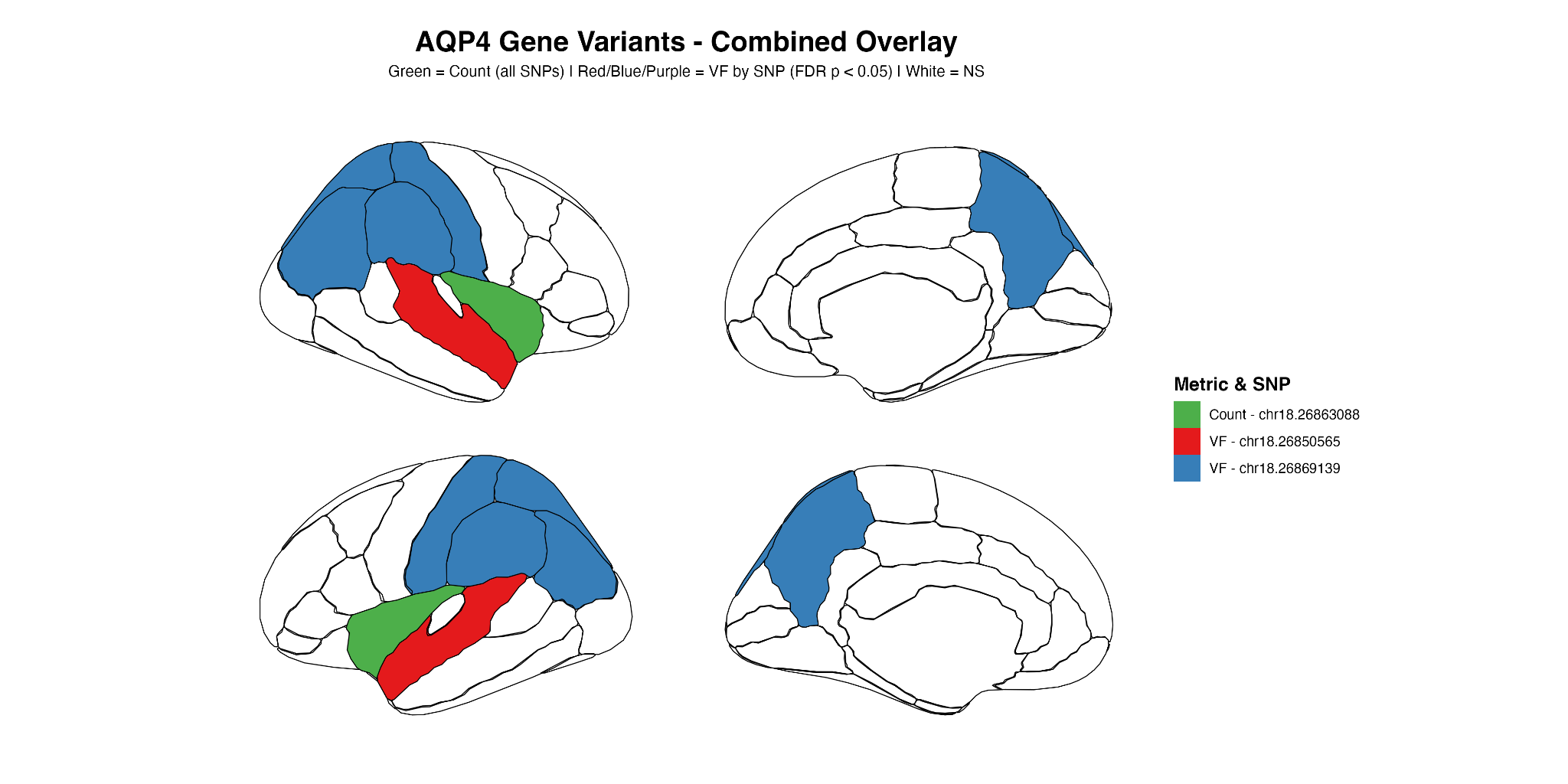
